# PD-L1^hi^ plasmablasts limit the T cell response during the acute phase of parasite infections

**DOI:** 10.1101/2020.06.10.142067

**Authors:** Melisa Gorosito Serrán, Facundo Fiocca Vernengo, Laura Almada, Cristian G Beccaria, Pablo F Canete, Jonathan Rocco Alegre, Jimena Tosello Boari, Maria Cecilia Ramello, Ellen Wehrens, Yeping Cai, Elina Zuniga I, Carolina L Montes, Eva V Acosta Rodriguez, Ian A. Cockburn, Carola G Vinuesa, Adriana Gruppi

**Affiliations:** Centro de Investigaciones en Bioquímica Clínica e Inmunología (CIBICI)-CONICET, Facultad de Ciencias Químicas, Universidad Nacional de Córdoba, X5000HUA Córdoba, Argentina; Department of Immunology and Infectious Disease, John Curtin School of Medical Research, Australian National University, Canberra, ACT, Australia; Division of Biological Sciences, University of California San Diego, La Jolla, San Diego, CA 92093, USA; China-Australia Centre for Personalised Immunology, Shanghai Renji Hospital, Shanghai Jiaotong University, Shanghai, China

**Keywords:** Plasmablasts, B cell, PD-L1 - Trypanosoma cruzi, Plasmodium, LCMV - immunosuppression, microorganism evasion

## Abstract

During infections with protozoan parasites or virus, T cell immunosuppression is generated simultaneously with a high B cell activation. Here, we show that in *T. cruzi* infection, all plasmablasts detected had higher surface expression of PD-L1, than other mononuclear cells. PD-L1^hi^ plasmablasts were induced *in vivo* in an antigen-specific manner and required help from Bcl-6^+^CD4^+^T cells. PD-L1^hi^ expression was not a characteristic of all antibody-secreting cells since plasma cells found during the chronic phase of infection express PD-L1 but at lower levels. PD-L1^hi^ plasmablasts were also present in mice infected with *Plasmodium* or with lymphocytic choriomeningitis virus, but not in mice with autoimmune disorders or immunized with T cell-dependent antigens. PD-L1^hi^ plasmablasts suppressed T cell response, via PD-L1, *in vitro* and *in vivo*. Thus, this study reveals that extrafollicular PD-L1^hi^ plasmablasts, which precede the germinal center (CG) response, are a suppressive population in infections that may influence T cell response.

**Brief summary:** Pathogens develop different strategies to settle in the host. We identified a plasmablats population induced by pathogens in acute infections which suppress T cell response.

## INTRODUCTION

Persistent replicating parasites and viruses can overcome the initial anti-microorganism response by suppressing innate and T cell response to allow the establishment of chronic infection. The mechanisms involved in this immunosuppression include different cell types and pathways, making it challenging to develop effective therapies. Immune suppression in chronic infections serves both to prevent inflammatory immune pathogenesis and to suppress protective effector immune responses ^1,2^. As such, *Trypanosoma cruzi* infection, also known as Chagas disease, has been a relevant model to understand the delicate balance between protective T cell immunity, immune suppression and parasite establishment.

Chagas disease presents an acute phase, both in mice and humans, characterized by a state of immunosuppression in which *T. cruzi* replicates extensively and induces immunomodulatory molecules that delay parasite-specific T cell effector responses ^3–6^. It has been reported that during this phase of *T. cruzi* infection the expression of PD-1, one of the members of the CD28/CTLA4 family with inhibitory capacity, increases on myocardium infiltrating CD4^+^ and CD8^+^ T cells ^7^ . The cross-linking of PD-1 with any of its two ligands, PD-L1 ^8^ or PD-L2 ^9^, inhibits the activation of T cells and the production of IL-2 and IFNγ.

Many studies have shown that other protozoan parasites such as *Plasmodium* spp. as well as bacteria and viruses also benefit from the PD-1/PD-L1 pathway to suppress and evade the host’s adaptive immunity ^10,11^. The activation of the PD-1/PD-L1 pathway would enable the establishment of microorganisms and chronicity ^12–14^. Accordingly, the blocking of PD-1/PD-L1 signals, in combination with therapeutic immunizations or therapies with cytokines, has been proposed as a strategy to improve the efficacy of vaccinations ^15^ and to revitalize exhausted T cells ^16^.

During infection with *T. cruzi*, blocking PD-1 and PD-L1 or deletion of the PD-1 gene results in a reduction in parasitemia and tissue parasitism, but also increased mortality due to an augmented cardiac inflammatory response ^7^ . These results reveal that the signaling pathway of PD-1/PD-L1 has an important role in the control of inflammation induced by *T. cruzi* and provide another perspective on the mechanisms of regulation in the pathogenesis of Chagas disease. However, PD-L1-expressing regulatory cells have not been fully identified in this context.

It has been reported that PD-L1^+^ regulatory plasmablasts have a critical role in suppressing immune T cell responses in cancer models ^17^ ; however, their role in chronic infection has not been described. Recently, we reported that Blimp1^f/f^CD23^Cre^ mice, which are deficient in plasmablasts, had a significant higher number of trypomastigotes in blood and splenic TNF^+^CD4^+^ T cells in comparison to infected CD23^Cre^ mice after *T. cruzi* infection ^18^, suggesting that plasmablasts and plasma cells are important for parasite replication control and also for the regulation of TNF-producing cells. Considering that the abscence of plasmablasts improves effector T cell responses, we explored whether PD-L1^+^ plasmablasts are present in *T. cruzi* infected mice and in other parasitic and chronic infections. Moreover, given the proximity of extrafollicular plasmablasts to T cell zones ^19^, we further asked whether PD-L1+ plasmablasts can regulate the T cell response via PD-1/PD-L1 pathway.

Here we show that, in *T. cruzi* infection, all plasmablasts detected had higher surface expression of PD-L1, than other B-lymphocyte lineage cells or other mononuclear cells. PD-L1^hi^ plasmablasts were induced *in vivo* in an antigen-specific manner and required help from Bcl-6^+^CD4^+^T cells. PD-L1^hi^ expression was observed in splenic plasmablasts detected during the acute phase of the infection but not in the bone marrow plasma cells found during the chronic phase of infection. PD-L1^hi^ plasmablasts were also present in mice infected with *Plasmodium* or with lymphocytic choriomeningitis virus (LCMV), but not in mice with autoimmune disorders or immunized with T cell-dependent antigens. We were also able to ascribe a suppressive role to these cells *in vitro*, and also *in vivo* in *T. cruzi* and *P. chaubadi* infections. In particular we found that PD-L1^hi^ plasmablasts suppressed IFNγ^+^TNF^+^PD1^+^CD4^+^ T cells via PD-L1. This study reveals that extrafollicular PD-L1^hi^ plasmablasts, which precede the germinal center (CG) response, are a suppressive population in infections that may influence T cell response. providing new insights that could be harnessed for cell targeting immune therapies.

## RESULTS

### The plasmablast response preceds GC formation and declines before CD4+ T cells peak

To understand the mechanisms by which plasmablasts regulate the CD4^+^ T cell response in experimental Chagas disease, we first investigated the kinetics of the plasmablast response. Compared with uninfected mice, flow cytometry analysis revealed that the number of splenic plasmablasts (CD3^−^B220^low^IgD^−^CD138^+^) increased prior to germinal center (GC) reaction, and peaked at 15-18 days post infection (Dpi) with *T. cruzi* (Fig.1a). The number of plasmablasts declined at 20 Dpi, whereas the number of GC B cells (CD19^+^CD3^−^CD138^−^B220^+^CD38^low^Bcl6^+^) remained high until 25 Dpi and decreased later. Interestingly, the peak of the CD4^+^ T cell response occurred only after the frequency and number of plasmablasts had decreased (Fig.1a)

**Fig. 1:**
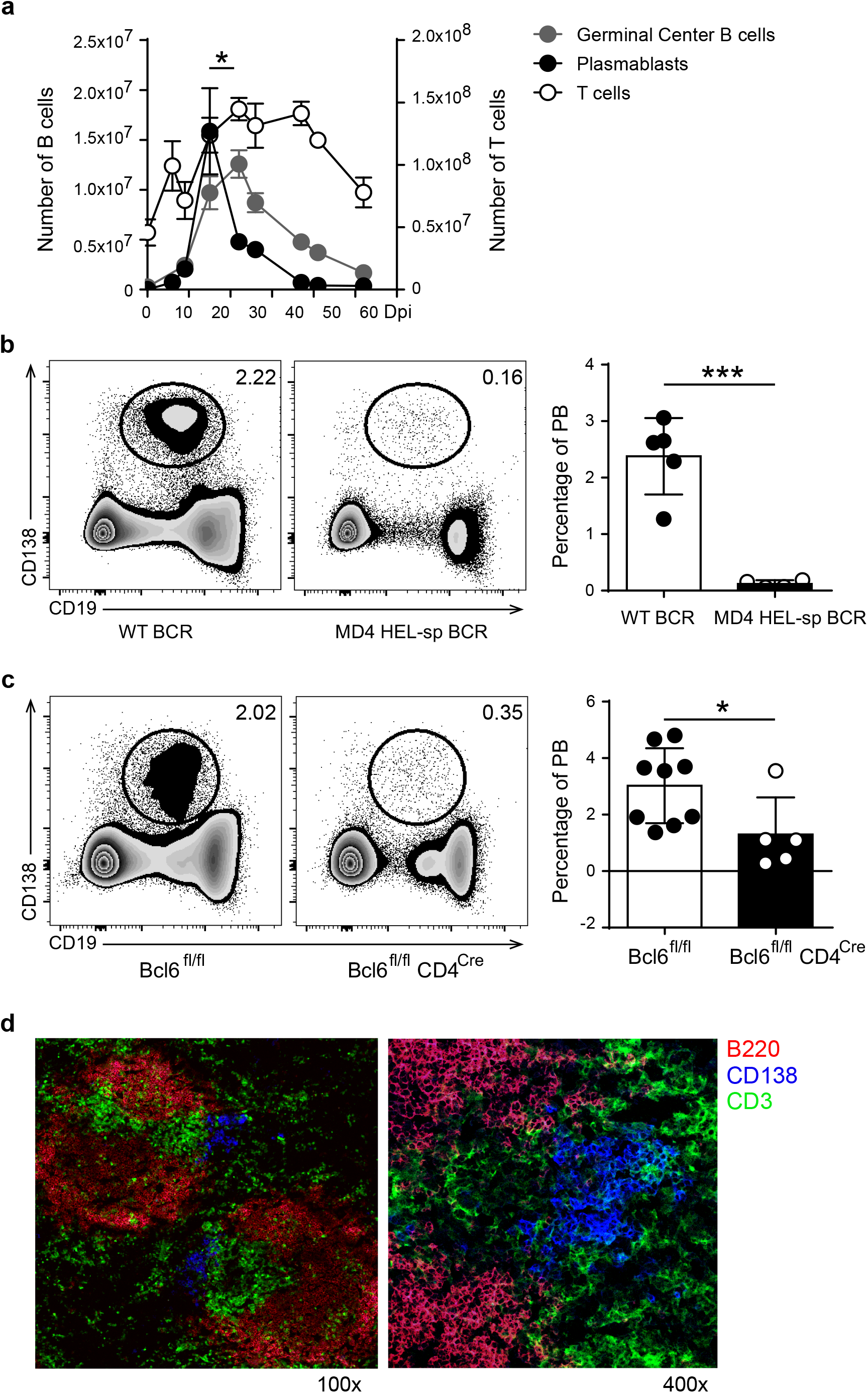
In *T. cruzi* infection plasmablast response was previous to GC reaction and was generated in an antigen specific manner and required Bcl-6^+^CD4^+^T cells collaboration. **a)** Kinetics of plasmablasts (PB, CD19^int^B220^low^CD138^hi^), Germinal Center (GC) B cells (CD19^+^CD3^−^CD138^−^ B220^+^CD38^low^Bcl6^+)^ and CD3^+^CD4^+^ T cells number in the spleen of *T. cruzi* infected mice. **b)** Representative flow cytometry plots and statistical analysis of the percentage of splenic PB (CD138^+^CD19^int^/^low^) from WT (WT BCR, n=5) and MD4 (HEL-sp BCR, n=8) mice at 14 Dpi with *T. cruzi*. Correlation between the percentage of PB and the percentage of IgHEL B cells (non-related BCR) in the spleen from infected MD4 (white circles) or WT (black circles) mice (***p < 0.001, two tailed t Test). **c)** Representative flow cytometry plots and statistical analysis of the percentage of splenic PB from WT (n=9) and Bcl6^fl/fl^CD4^Cre^ (Bcl-6^neg^CD4+ T cells, n=5) mice at 14 Dpi with *T. cruzi*. (**p < 0.01, two tailed t Test) **d)** Representative Immunofluorescence of spleen sections (7um) from *T. cruzi*-infected C57BL/6 mice obtained at 14 Dpi, stained with anti-B220 (red), anti-CD3 (green) and anti-CD138 (blue). Magnification: x100 (left) and x400 (right) (n = 9).

### Plasmablasts are generated in an antigen specific and Tfh-dependent manner

To understand in more depth the origin and characteristics of plasmablasts in *T. cruzi* infection, we infected MD4 mice (whose B cells express a transgenic BCRs specific for a *T. cruzi* non-related protein, HEL) and their non-transgenic controls (WT-BCR, whose B cells express a wild-type BCR repertoire). Infection of MD4 mice showed that plasmablasts were generated in an antigen-specific manner, since plasmablasts were almost absence from infected MD4 mice compared to wild type mice around the peak of the plasmblasts responses at 16 Dpi (Fig. 1b, left and right graphs).

Since extrafollicular (EF) antibody responses requires Bcl-6 expression by T cells ^20^, we next investigated whether plasmablast induction and/or survival required the collaboration of Bcl-6^+^CD4^+^T cells. For this, we took advantage of Bcl6^fl/fl^CD4^Cre^ mice that lack Tfh and CXCR5^neg^Bcl-6+CD4+ T cells. While Cre-negative littermates develop a normal Bcl-6^+^CD4^+^T cell response ^21^, *T. cruzi* infected Bcl6^fl/fl^CD4^Cre^ mice had a significant decrease in the frequency of plasmablasts (Fig. 1c, right graph), indicating the plasmablast response is critically dependent on Bcl-6^+^CD4^+^ T cells.

Immunofluorescence analysis of the spleen from *T. cruzi* infected mice (Fig. 1d) identified CD138^+^ cells extending beyond the classical sites of extrafollicular plasmablast growth (Fig. 1d, CD138+ cells, blue) into the T zone areas (Fig. 1d, CD3^+^ cells, green), a finding consistent with our previous reports ^19,22^. The results suggested that in *T cruzi* infection, plasmablasts were induced in a BCR-dependent manner, required Bcl-6^+^CD4^+^ T cell collaboration and were located at extrafollicular sites in close proximity to T cells.

### Plasmablasts from T. cruzi infected mice express high levels of PD-L1

To identify if plasmablasts express inhibitory molecules able to modulate T cell response, we evaluated the expression of PD-L1 and PD-L2 ^23^ on plasmablasts from *T. cruzi* infected mice. In comparison with B and non-B cells, plasmablasts from *T. cruzi* infected mice, evaluated at different Dpi, expressed the highest levels of PD-L1 (determined as MFI, Fig. 2a). Remarkably, all of the plasmablasts detected during the acute phase of *T. cruzi* infection expressed high levels of PD-L1 (Fig. 2a). As a consequence of the infection, B cells and plasmablasts also expressed PD-L2, but at similar levels (Fig. 2a, see histograms and bar graph). PD-L1^hi^ plasmablasts were absent in the spleen of infected mice during the chronic phase of the infection (130 Dpi, data not shown). Interestingly, while most plasmablasts analyzed during the acute phase were PD-L1^+^, only ~ 50% of the bone marrow plasma cells detected in the chronic phase (130 Dpi) expressed PD-L1 (Fig.2b, histograms). In addition, PD-L1 expression on plasma cells detected in the chronic infection was significantly lower respect to PD-L1 expression on plasmablasts present at 15 Dpi (Fig. 2b, bar graph).

**Fig. 2.**
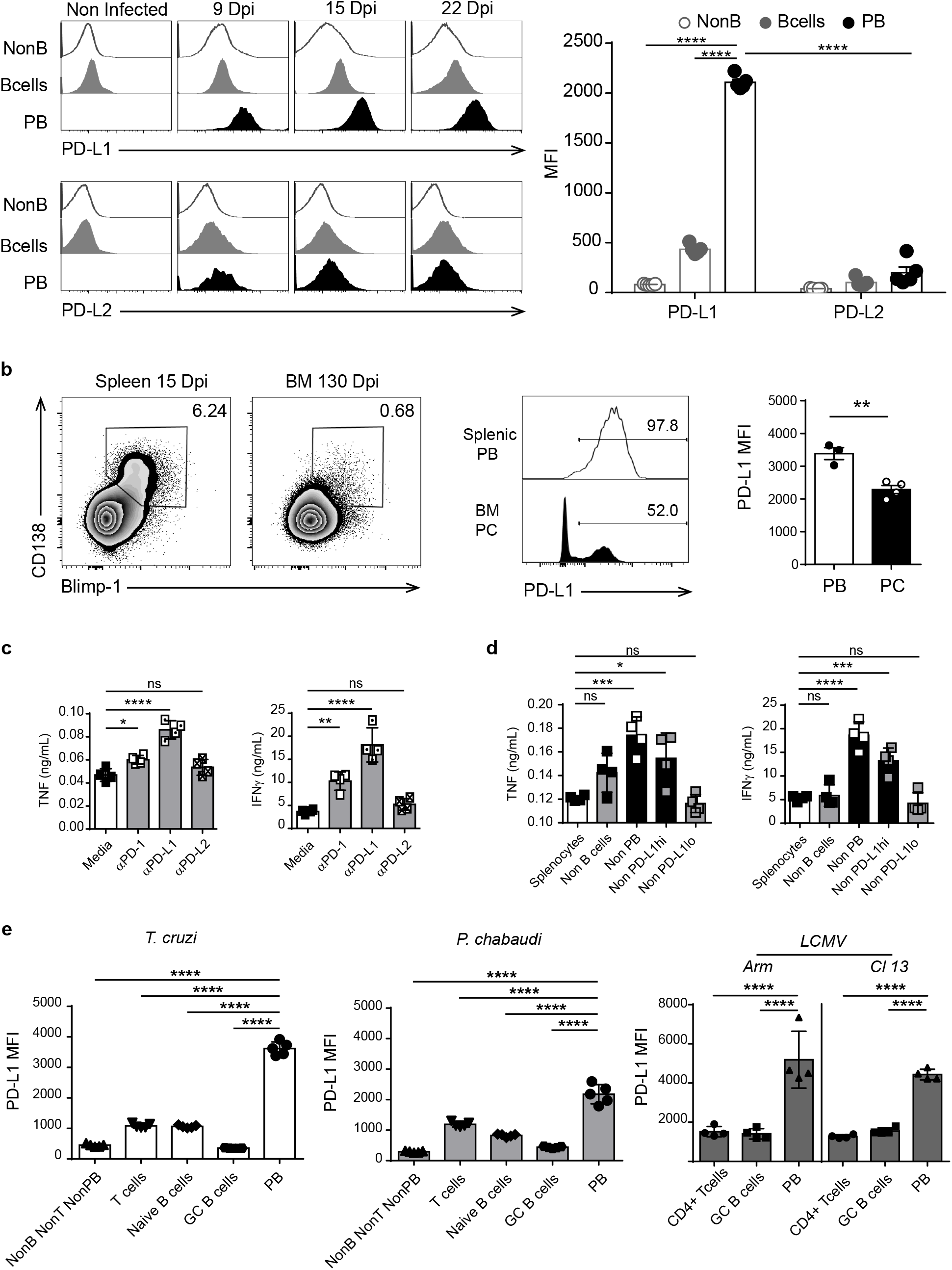
Plasmablasts from *T. cruzi* infected mice expressed high levels of PD-L1. a) Representative histograms of PD-L1 or PD-L2 expression on splenic non-B cells (CD19^−^CD138^−^B220^−^), B cells (CD19^+^CD3^−^CD138^−^B220^+^) from non-infected mice and *T. cruzi*-infected C57BL/6 mice and on plasmablasts (PB) (CD19^int^B220^low^CD138^hi^) from *T. cruzi*-infected C57BL/6 mice evaluated at 9, 15 and 22 Dpi. Statistical analysis of mean fluorescence intensity (MFI) of PD-L1 or PD-L2 on the mentioned cells analyzed at 15 Dpi. **b)** Representative plots of CD138 vs Blimp-1 of live single lymphocytes (gated on IgD^−^IgM^−^CD11b^−^CD24^−^) from the spleen and bone marrow (BM) obtained at 15 Dpi and 130 Dpi with *T. cruzi*, respectively. Representative histograms of PD-L1 on gated CD138^+^Blimp-1^+^ cells. Numbers indicate the percentage of PD-L1^+^ cells. Bar graph shows statistical analysis of MFI of PD-L1^+^ on splenic PB and BM plasma cells (PC). **c)** 10^5^ Splenocytes from *T. cruzi* infected mice obtained at 14 Dpi were incubated with *T. cruzi* antigens and cultures for 48h with media, anti-PD-1, anti-PD-L1 or anti-PD-L2. TNF and IFNγ concentrations were determined by Elisa in the culture supernatant. **d)** 1× 10^5^ Non-B (CD19^neg^CD138^neg^) cells in a suspension of total splenocytes (non-depleted) from *T. cruzi* infected mice obtained at 14 Dpi or depleted of B cells (Non B cells), plasmablasts (Non PB), PD-L1^hi^ cells (Non PD-L1^hi^) or PD-L1^low^ (Non PD-L1^lo^) cells were cultured with *T. cruzi* antigens for 48h. TNF and IFN✉ concentrations were determined by Elisa in the culture supernatant. **e)**Statistical analysis of MFI of PD-L1 on Non-B non-T non-PB cells (CD3^−^CD19^−^B220^−^CD138^−^), T cells (CD3^+^CD19^−^B220^−^), naïve B cells (CD19^+^CD3^−^CD138^−^B220^+^IgD^+^CD38^+^Bcl6^−^), GC B cells (CD19^+^CD3^−^CD138^−^B220^+^CD38^low^Bcl6^+^) and PB (CD19 ^int^CD138^+^) from the spleen of *T. cruzi* infected mice obtained at 14 Dpi, or from the spleen of *P. chabaudi* infected mice obtained at 10 Dpi; or CD4+ T cells, GC B cells of PB from the spleen of Arm or Cl13 LCMV infected mice obtained at 9 Dpi.*p < 0.05, ** p < 0.01, ***p < 0.001, ****p < 0.0001 one -way ANOVA, Bonferroni post-test

To test the role of PD-L1 and PD-L2 on TNF- and IFNγ–producing cells from *T cruzi* infected mice, splenocytes stimulated with *T. cruzi* antigens were cultured with medium or with anti-PD-1, anti-PD-L1 or anti-PD-L2 and the concentration of cytokines was evaluated in the culture supernatant. The blockade of PD-1 and PD-L1 signaling significantly increased TNF and IFNγ concentration in the culture supernatants (Fig. 2c). No significant changes were detected in the concentration of cytokines in the supernatants of splenocytes cultured with anti-PD-L2 compared to splenocytes cultured with medium (Fig. 2c) suggesting that the PD-L2 pathway was not involved in the *in vitro* regulation of cytokine-producing cells from *T. cruzi* infected mice.

To identify the B-lineage cell population responsible for limiting cytokine-producing T cells, splenocytes from *T. cruzi* infected mice obtained at 14 Dpi were depleted of plasmablasts or B cells, by cell sorting (Supplementary Fig.1a-c). Later, the cells were cultured with parasite-antigens and cytokine concentration was determined in culture supernatants. Only plasmablast depletion significantly increased the concentration of TNF and IFNγ in the culture supernatant (Fig. 2d). Depletion of PD-L1^hi^ cells, but not of PD-L1^low^ cells from splenocytes also lead to an increase in the concentration of cytokines in the supernatant (Fig. 2d). Given that depletion of PD-L1^hi^ cells concomitantly leads to plasmablasts elimination (Supplementary Fig. 1d and e), these data suggest that plasmablasts with high PD-L1 expression inhibit cytokine-producing cells *in vitro*.

We next investigated whether PD-L1^hi^ plasmablasts are unique to *T. cruzi* infected mice or if this population is present in other persistent infections. Analysis of the spleens of mice infected with *P. chabaudi* or with LCMV (either Armstrong –ARM- or clone 13 –Cl13-isolates) analyzed after 10 and 9 Dpi, respectively, showed similar frequencies of PD-L1^+^plasmablasts in the different infection conditions (Supplementary Fig. 2). Similar to our findings with *T. cruzi*, the plasmablasts from *P. chabaudi* and from ARM- or Cl13-LCMV infected mice also expressed high levels of PD-L1 in comparison to T cells and GC B cells from the same mice (Fig. 2e).

### PD-L1^hi^ plasmablasts are not detected in autoimmune nor in immunized mice

We next investigated whether PD-L1^hi^ plasmablasts occur in autoimmune disease or after immunization with non-microbial protein antigens. Mice known to develop spontaneous autoimmunity such as *Fas*^*lpr*^, *sanroque* and *Trex1*-deficient mice had varying frequencies of splenic plasmablasts (Fig. 3a). Regardless of the frequency of plasmablasts present in the autoimmune mice, PD-L1 expression on splenic plasmablasts from autoimmune mice was significantly lower than on plasmablasts in mice infected with *T. cruzi* (Fig. 3b), although they remained positive compared to the isotype control. To test if protein antigen-immunization that induces EF plasmablasts triggers PD-L1 expression on plasmablasts, we used the SwHEL cell transfer model ^24^ . HEL-specific B cells from SwHEL mice were adoptively transferred into C57BL/6 mice which were then immunized iv with sheep red blood cells (SRBC) alone (immunization control) or SRBC conjugated to HEL2x (Paus et al., 2006). PD-L1 expression at the peak of the EF HEL-specific plasmablast response (day 6) was significantly lower than that from plasmablasts at the peak of acute *T. cruzi* infection (14 Dpi) (Fig. 3c, histograms). Importantly, HEL-specific plasmablasts had a similar PD-L1 expression to non-HEL binding plasmablasts and to plasmablasts generated by SRBC alone (Fig. 3d). Overall, the results suggest that BCR binding on B cells was not sufficient for high PD-L1 expression on plasmablasts.

**Fig. 3.**
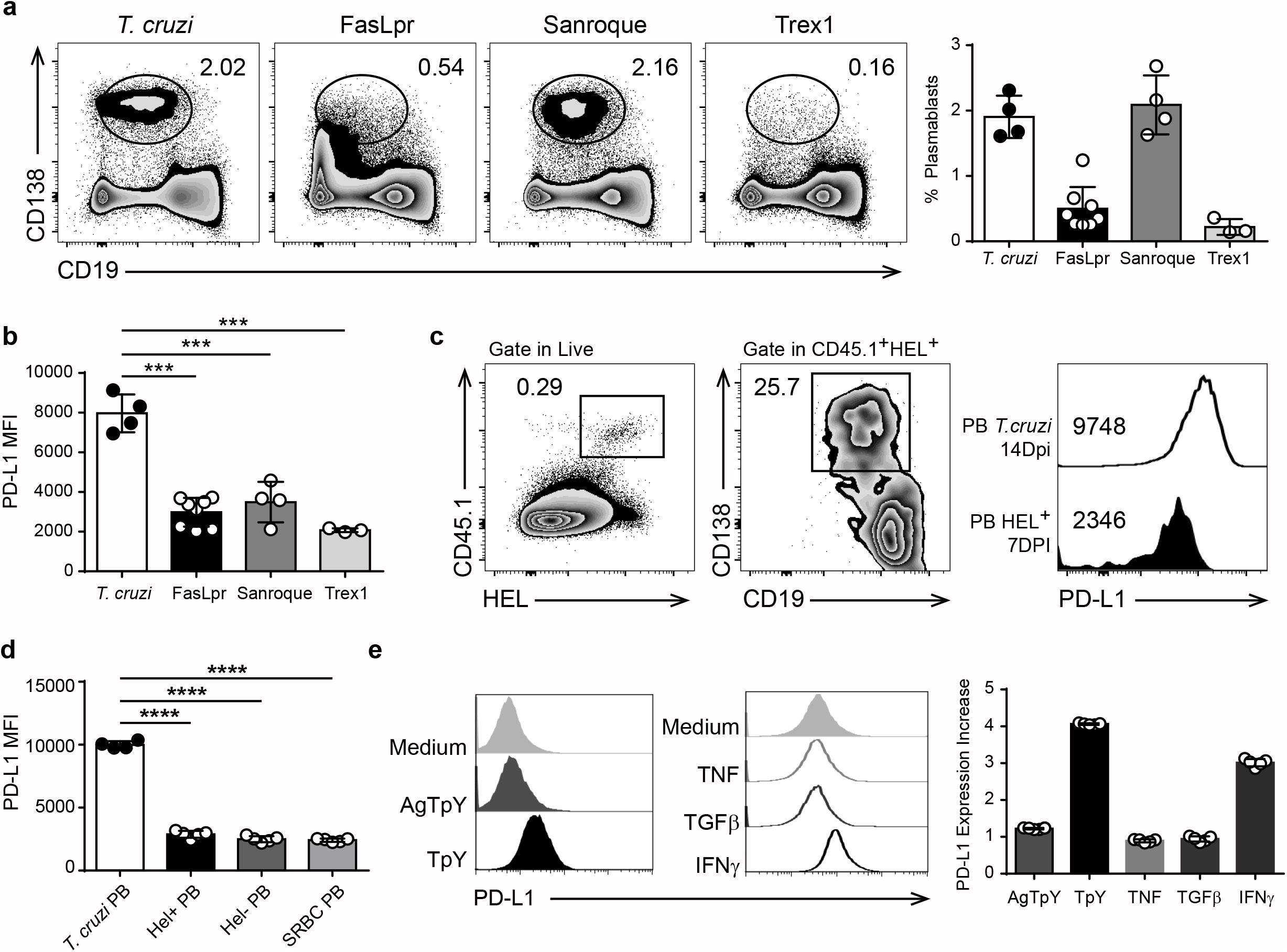
PD-L1^hi^ plasmablasts were present in infected but not in autoimmune mice. a) Representative flow cytometry plots and statistical analysis of the percentage of splenic plasmablasts from *T. cruzi-infected* and from *FasLpr, Sanroque* and Trex1 mice. C57BL6 mice (n=4) were infected with *T. cruzi* Y strain and the spleens were obtained at 14 Dpi. Spleens from *FasLpr (n=7), Sanroque (n=4)* and Trex1 *(n=3)* mice were obtained at 10-12 weeks of age. Splenocytes were stained with anti-B220, anti-CD138 and anti-PD-L1. Data are representative of two independent experiments. Statistical analysis of PD-L1 expression (MFI) on plasmablasts (PB) from the spleen of *T cruzi-*infected mice and FasLpr, Sanroque and Trex1 mice. ***p < 0.001one-way ANOVA, Bonferroni post-test. **c)** Representative flow cytometry plots of the percentage of SWHEL-specific B cells identified as CD45.1^+^HEL^+^ cells and the percentage of SWHEL-specific plasmablasts CD138^+^CD19^low^ in CD45.2 mice transferred with 30.000 HEL-specific B cells and immunized with 2×10^8^ SRBC conjugated with HEL. Histograms show PD-L1 expression on plasmablasts from 14 Dpi-*T. cruzi* infected mice and SWHEL-specific plasmablasts (PB) evaluated at 7 days post immunization (DPI). **d)** Statistical analysis of PD-L1 expression (MFI) on plasmablasts from *T. cruzi* infected mice, SWHEL-specific (HEL^+^PB) and –non-specific (HEL^neg^PB) plasmablasts from CD45.2 mice transferred with 30.000 HEL-specific B cells and immunized with 2×10^8^ SRBC conjugated with HEL, and on plasmablasts from mice immunized with SRBC (7 DPI). ****p < 0.0001 one -way ANOVA, Bonferroni post-test. **e)** Representative histograms of PD-L1 expression on B cells cultures with medium, *T. cruzi* trypomastigotes antigens (AgTpY), live *T. cruzi* trypomastigotes (TpY), TNF, TGFβ and IFNγ. Bar graph shows statistical analysis of PD-L1 expression increase on B cells cultured with the mentioned stimulus respect to medium.

Since PD-L1^hi^ expression on plasmablasts was observed in infected mice, we hypothesize that a strong inflammatory environment or the pathogens favors the high expression of PD-L1 on plasmablasts. To test this, purified B cells from uninfected C57BL/6 mice were cultured in the presence or absence of *T. cruzi* antigens, live trypomastigotes or cytokines, and PD-L1 expression was evaluated after 24h. In comparison to B cells cultured with medium alone, PD-L1 upregulation was observed when B cells were cultured with live trypomastigotes or with recombinant IFN✉ but not with *T. cruzi* antigens, TNF or TGF✉ (Fig. 3e). However, PD-L1^hi^ plasmablasts were present in IFN✉−/− mice infected with *T. cruzi* (data not shown), indicating that this cytokine may be redundant with other cytokines for mediating PD-L1 upregulation on plasmablasts *in vivo*. Together, these results suggest that live parasites are required to induce maximal PD-L1 expression, possibly through by inducing a combination of pro-inflammatory cytokines.

### Plasmablasts regulate IFNγ and TNF-producing cells via PD-L1

To test whether plasmablasts suppress T cell responses *in vivo* in a PD-L1 dependent manner, we set up mixed bone marrow chimeras in which only plasmablasts lacked PD-L1 expression. This was achived by transferring 80% *Blimp1*^f/f^CD23^Cre^: 20% *Pdl1*^−/−^ bone marrow cells into sub-lethally irradiated *Rag*1^−/−^ recipients. As controls, we used C57BL/6 (WT) and Blimp1^f/f^CD23^Cre^ (plasmablast deficient) chimeric mice. At the peak of the plasmablast response to *T. cruzi* (15-18 Dpi), the frequency and number of PD-1^+^ T cells and PD-1^+^IFNγ^+^TNF^+^ CD4^+^ T cells were evaluated. Flow cytometry analysis showed that, in *T. cruzi*-infected mice lacking PD-L1 expression on plasmablasts, there was a significant increase in the percentages and absolute numbers of PD-1^+^CD4^+^ T cells and IFNγ^+^TNF^+^ PD-1^+^CD4^+^ T cells (Fig. 4a). In chimera mice deficient in the whole plasmablast population only the frequency of PD-1^+^CD4^+^ T cells was increased in comparison to control chimera mice (Fig.4a, white bars).

**Fig. 4.**
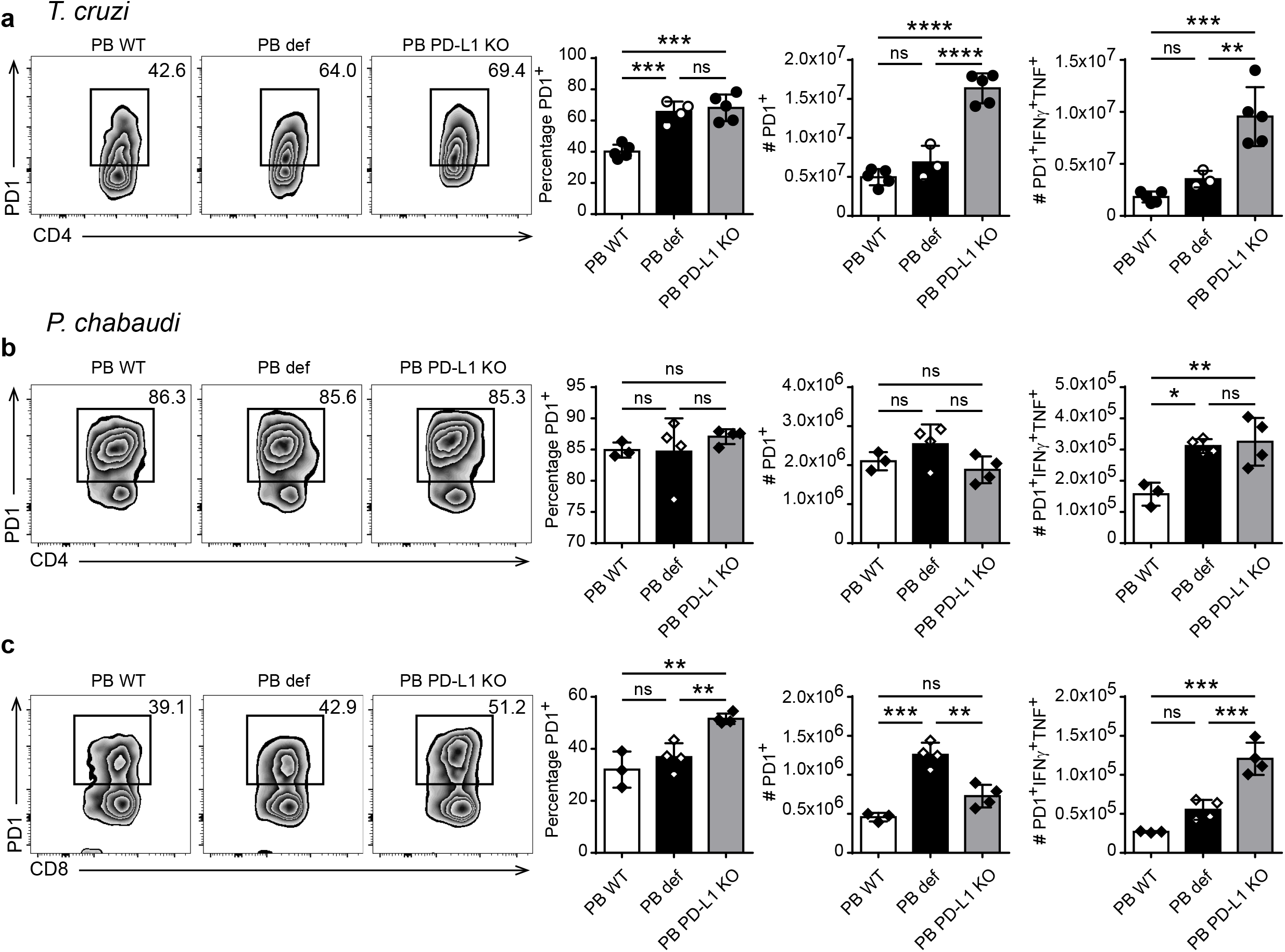
PD-L1 ^hi^ Plasmablasts were able to regulate IFN✉ and TNF-producing cells via PD-L1. **a)** Representative flow cytometry plots of PD1^+^ vs CD4^+^ cells and statistical analysis of the percentage of splenic effector PD1^+^CD4^+^ T cells and of the number of PD1^+^CD4^+^ T cells and IFNγ^+^TNF^+^PD1^+^CD4^+^ T cells from *T. cruzi-*infected chimeras. Recipient RAG1 KO mice (n=13) were sublethally irradiated and reconstituted via iv injection with 10×10^6^ bone marrow cells from C57BL/6 mice (WT) (n=5), Blimp^fl/fl^-iCre^CD23^ (PB def) (n=3), or a combination of an 80:20 mixture of 10×10^6^ BM cells from Blimp^fl/fl^-iCre^CD23^ plus PD-L1 KO donor mice (n=5). After 10 weeks of reconstitution, mice were infected with *T. cruzi* trypomastigotes Y strain and the spleens were obtained at 14 Dpi. Splenocytes were stained for viability and with anti-CD3, anti-CD4, anti-CD8, anti-CD44, anti-Foxp3, anti-PD1, anti-CXCR5, anti-IFNγ and anti-TNF. Gating of effector CD4+ T cells was performed: CD3^+^CD4^+^CD44^+^Foxp3^−^CXCR5^−^. **b)** Representative flow cytometry plots of PD1^+^ vs CD4^+^ cells and statistical analysis of the percentage of splenic effector PD1^+^CD4^+^ T cells and of the number of PD1^+^CD4^+^ T cells and IFNγ^+^TNF^+^PD1^+^CD4^+^ T cells and **c)** CD8+ T cells from *P. chabaudi-infected* chimeras. Recipient RAG1 KO mice (n=11) were sublethally irradiated and reconstituted via iv injection with 10 × 10^6^ bone marrow cells from a combination of an 80:20 mixture of 10.10^6^ BM cells from Blimp^fl/fl^-iCre^CD23^ plus C57BL/6 donor mice (WT) (n=3), Blimp^fl/fl^-iCre^CD23^ (PB def) (n=4), or a combination of an 80:20 mixture of 10×10^6^ BM cells from Blimp^fl/fl^-iCre^CD23^ plus PD-L1 KO donor mice (n=4). After 10 weeks of reconstitution, mice were infected with *P chabaudi* and the spleens were obtained at 10 Dpi. Splenocytes were stained for viability and with anti-CD3, anti-CD4, anti-CD8, anti-CD44, anti-Foxp3, anti-PD1, anti-CXCR5 anti-IFNγ and anti-TNF. Gating of effector CD4+ T cells was performed: CD3^+^CD4^+^CD44^+^Foxp3^−^CXCR5^−^. Gating of effector CD8+ T cells was performed: CD3^+^CD8^+^CD44^+^. Experiment was performed twice. *p < 0.05, **p < 0.01, ***p < 0.001, ****p < 0.0001 one-way ANOVA, Bonferroni post-test.

To extend our findings beyond *T. cruzi* infection, we further evaluated whether PD-L1-mediated plasmablast immunosuppression occurred during malaria. Accordingly, the chimeras described above were infected with *P. chaubadi* and evaluated at 10 Dpi, which corresponds to the peak of PD-L1^hi^ plasmablast expansion in this infection. We observed a significant increase in the number of IFNγ^+^TNF^+^PD-1^+^CD4^+^ T cells from the spleens of mice that are unable to generate plasmablasts (Fig. 4b, black bar) or that specifically lack the expression of PD-L1 on plasmablasts (Fig. 4b, grey bar) compared to WT chimeras. In addition, we observed that there was a significant increase in the percentage of PD-1^+^CD8^+^ T cells and in the number of IFNγ^+^TNF^+^PD-1^+^CD8^+^ T cell population in chimeras with PD-L1KO plasmablasts (Fig.4c). Together, these results suggest that in *T. cruzi* infection and malaria, PD-L^hi^ plasmablasts are able to suppress IFN✉ and TNF-producing PD-1^+^ T cells.

## Discussion

Our current understanding of humoral responses has been largely derived from immunizations with model antigens. In the last decade, murine models of infection that also present with a significant inflammatory component have allowed the identification of new cell populations that have fundamental roles in the induction of the B cell response ^25,26^, the discovery of novel sites where B cells are activated ^27^ and identification of distinct regulatory B cell functions ^22,28^. Here, we describe plasmablasts with very high expression of PD-L1 that were detected in infected mice but absent from mice with autoimmunity or immunized with T-dependent model antigens. Our findings provide the first evidence that most of the plasmablasts in *T. cruzi*, *Plasmodium* or LCMV infections express high levels of PD-L1, in comparison to naïve and GC B cells, T cells or other lymphoid cells. Interestingly, CD138^+^B220^low^Blimp-1^hi^ cells present in bone marrow at the chronic phase of *T. cruzi* infection did not exhibit high expression of PD-L1, suggesting that PD-L1^hi^ expression is not a characteristic of all antibody-secreting cells. PD-L1 expression was highest in an antibody-secreting cell population present in the acute phase of the infection, when there is a strong inflammation and the effector response of T cells begins to develop ^29^.

All the plasmablasts detected during the acute phase of *T. cruzi* infection are PD-L1^hi^ and are extrafollicular since the CD138^+^B220^low^ cells, located outside the follicle, are the only antibody-secreting cells expressing CD138 at this time of infection ^19^ and Figure 1). The appearance of PD-L1^hi^ plasmablasts precedes GC formation and, as previously reported for extrafollicular response ^20^, Bcl-6 expression in T cells has been shown to be essential for their optimal induction/survival. Consistent with this, very few PD-L1^hi^B220^low^ cells were detected in the spleens of infected mice lacking Bcl-6 expression in CD4+T cells. Additionally, despite the prevailing views that during the acute phase of *T. cruzi* infection most of B cells lack parasite specificity suggesting non-antigen specific B cell activation ^30^, PD-L1^hi^ plasmablasts required parasite-specific BCRs. Indeed, mice whose B cells expressed BCRs non-specific for a *T. cruzi* exhibited very low number of plasmablasts.

PD-L1 high expression on antibody-secreting cells was previously reported on intestinal lamina propria IgA+ plasma cells, which are the major PD-L1 expressing cells in that tissue. PD-L1 expression levels on intestinal IgA plasma cells are higher than that on IgG plasma cells in peripheral lymphoid tissues ^31^. Probably, the high expression of PD-L1 in the intestinal IgA plasma cells is triggered by the microbiota, since we observed that microorganisms (in this report, *T. cruzi*) per se and/or inflammatory cytokines can induce high PD-L1 expression on B cells. PD-L1 expression on plasmablast was also described in different mouse prostate cancer models under Oxaliplatin treatment ^17^. This therapy induces tumor-infiltrating plasmocytes which express IgA, IL-10 and PD-L1. Elimination of these cells, which also infiltrate human-therapy-resistant prostate cancer, allows cytotoxic T cell-dependent eradication of oxaliplatin-treated tumors ^17^, suggesting a suppressive function of this population.

Considering PD-L1 plays a major role in suppressing the adaptive arm of immune system, PD-L1^hi^ plasmablasts emerge as a population with regulatory capacity. The majority of studies on Breg cells have focused on the predisposition of immature and mature B cells to produce IL-10 and to suppress a range of autoimmune or allergic conditions in both mice and man ^32^. Little information is available about the regulatory function and mechanisms of action of antibody-secreting cells. In this study, we have identified that plasmablasts from *T. cruzi* or *Plasmodium* infected mice exert their regulatory function through PD-L1. When, splenocytes from *T. cruzi* infected mice were depleted from PD-L1^hi^ cells or plasmablasts, and later cultured with *T. cruzi* antigens, we observed a high increase in the levels of TNF and IFN✉ in the culture supernatant, respect to the culture of total splenocytes. This result indicates that the presence of PD-L1^hi^ plasmablasts controls/modulates effector cell response. The *in vitro* studies were supported by chimeric mice in which absence of PD-L1 on plasmablasts led to elevated frequency of PD-1^+^IFNγ-^+^TNF^+^-producing T cells, suggesting that PD-L1 expression on plasmablasts can control the inflammatory response.

It has been described that human and mouse glioblastoma associated B cells that overexpress PD-L1 also have immunosuppressive activity directed towards activated CD8+ T cells ^33^. B cells located in the tumor microenvironment but not circulating or from lymph nodes express high levels of PD-L1 but the PD-L1 expression apparently does not appear to depend on the inflammatory environment instead tumor myeloid-derived suppressive cells deliver microvesicles transporting PD-L1, which is taken up by tumoral B cells.

Plasmablasts also express PD-L2, another molecule that interacts with PD-1 and suppresses T cell proliferation and cytokine release ^9^. However, we observed that PD-L2 was not mediating effector cell suppression since PD-L2 blockage in cell culture did not modified cytokine production.

The data obtained in this work provide information that allows us to partially explain why during infections with protozoan parasites, such as *T. cruzi* and *Plasmodium*, T cell immunosuppression is generated simultaneously with a high B cell activation. The acute phase of Chagas’ disease in mice and human is marked by a state of immunosuppression, which coexist with polyclonal B cell activation, in which *T. cruzi* replicates extensively and induces immunomodulatory molecules that delay parasite-specific responses mediated by effector T cells ^34^. Also, it has been reported that in the acute phase of infection with *P. chabaudi* the composition of the B cell compartment is altered with vigorous extrafollicular growth of plasmablasts and germinal center formation. In *Plasmodium* infection, extrafollicular foci of plasmablasts are visible from day 4 initiating a very strong plasma cell response. By day 10, plasma cells are localised in the periarteriolar region of the white pulp and form clusters occupying part of the area normally filled by T cells. During this phase of infection, as in *T. cruzi* infection, there is a delayed GC response and a significant reduction of the humoral response to T-dependent antigens ^35^. Interestingly, we also observed that PD-L1^neg^ plasmablast functioned as an enhancers of T cell response (Fig.4a and 4c) since in infected-chimera mice the frequency of PD-1^+^IFNγ-^+^ TNF^+^-producing T cells was higher than in mice without plasmablast. The results suggest that if the PD-1/PD-L1 pathway is not operational the plasmablasts could function as antigen presenting cells ^36^.

Last, we observed that mice infected with LCMV ARM or Cl13 exhibited high PD-L1 expression in plasmablasts at 9 dpi. Infection of mice with LCMV ARM and Cl13 has served as a powerful model system to study immune responses during acute and chronic viral infections, respectively ^2^ . In particular, LCMV Cl13 induces higher and more sustained PD-1 upregulation in T cells, and PD-1/PD-L1 interaction significantly contributes to T cell suppression and viral persistence ^37^ . Interestingly, LCMV infection also triggers polyclonal B cell activation ^38,39^. Given that virus-specific CD4 and CD8 T cells from LCMV Cl13 (but not ARM) infected mice express high levels of PD-1 ^37^, it is tempting to speculate that, as we show for *T.cruzi* and *P. chabaudi* infections, PD-L1^hi^ plasmablasts may also contribute to suppressing antiviral T cells during chronic LCMV infection. It should be noted that although PD-L1^hi^ plasmablasts are also induced after LCMV ARM acute infection, they are unlikely to exert a significant regulatory function in this setting, as virus-specific T cells from ARM-infected mice exhibit low levels of PD-1 expression ^37^.

In summary, our results have unveiled a new immunoregulatory pathway in the context of infections, which could provide new target for the rational design of new therapeutic treatments aimed at enhancing protective T cell responses during Chagas disease, Malaria and potentially other chronic infections.

## METHODS

### Mice

C57BL/6, and B6.129S7-Ifngtm1Ts/J (IFNγKO) mice were initially obtained from The Jackson Laboratories (USA). PD-L1 KO mice were provided by Dr Halina Offner, Oregon Health & Science University. These mice were housed and bred in the Animal Facility of the CIBICI-CONICET, FCQ-UNC.

Mice from the Australian National University Bioscience Services were maintained in specific pathogen–free conditions and had access to food and water ad libitum. Mice used in the experiments conducted at the John Curtin School of Medical Research (JCSMR) were:

HEL-IgM-BCR transgenic mice (MD4, knock in for a transgenic BCR specific for HEL, a *T. cruzi* non-related protein) were used. As controls of MD4 mice we used their littermates whose genotype was negative for the knock in and therefore their B cells express a complete BCR repertoire

SW_HEL_ CD45.1 mice which carry a Vk10k light chain transgene and a knocked in VH10 Ig heavy chain in place of the JH segments of the endogenous IgH gene that encode a high-affinity antibody for HEL, were obtained from the laboratory of R. Brink (Garvan Institute, Sydney, New South Wales, Australia).

Blimp^fl/fl^-CD23iCre mice were obtained from crossing of females Blimp^fl/fl^ -CD23iCre-negative and males Blimp^fl/fl^-CD23icre-positive. Blimp^fl/fl^-iCre^CD23^ mice are plasmablast/plasma cell-deficient mice. iCre-negative littermates were used as controls (plasmablast/plasma cell sufficient).

Bcl6^fl/fl^ mice were mated to CD4-cre mice to generate Bcl6^fl/fl^ Cre^CD4^ mice ^21^. Those mice expressing Cre lack Tfh, while T lymphocytes littermates Cre-negative will present a functional Bcl6 molecule and develop a normal Tfh response.

Mice experiments conducted at the CIBICI were approved by and performed in accordance with the guidelines of the Institutional Animal Care and Use Committee of the FCQ-UNC (Approval Number HCD 1525/14). Mice experiments conducted at JCSMR were approved by the Animal Experimentation Ethics Committee (Australian National University protocol numner 2016/17). Male and female mice were used at age-matched (8-12 weeks-old) and housed with a 12-h light-dark cycle.

Mice experiments conducted at the Division of Molecular Biology, Department of Biological Sciences, University of California, San Diego, were handled conformed to the requirements of the National Institutes of Health and the Institutional Animal Care and Use Guidelines of University of California, San Diego.

### Pathogens and Experimental Infections

*T. cruzi* infection: mice were infected intraperitoneally with 1 × 10^4^ trypomastigotes of *T. cruzi* Y-Br strain ^40^ diluted in a sterile solution of 1% glucose in PBS ^22^. Uninfected littermates were injected with 1% glucose in PBS and processed in parallel. Parasitemia was monitored by counting the number of viable trypomastigotes in blood after lysis with a 0.87% ammonium chloride buffer. Spleen and lymph nodes were collected at different Dpi for immune response analysis.

*Plasmodium chabaudi* infection: mice were infected intravenously (iv) with 10000 parasites of *P. chabaudi* (strain AS) obtained from a mouse infected 8 days previously from a frozen stock. The spleens of *P chabaudi* infected mice were obtained at 10 Dpi. LCMV infection: mice were infected with 2 × 10^6^ plaque-forming units (pfu) of LCMV Arm (ARM) or LCMV clone 13 (Cl13) iv via tail vein. Viruses were propagated on BHK cells and quantified by plaque assay performed on Vero cells ^41^. The spleens of LCMV infected mice were obtained at 9 Dpi.

### Antigen preparation

*T. cruzi* parasites (Y-Br strain) were cultured in NIH3T3 mouse fibroblasts and were collected as described ^40^. Trypomastigotes were washed twice in sterile PBS (37ºC) and were resuspended in 500 ul of PBS (4°C) and sonicated in ice-cold water for 10 minutes. Aliquots were stored at −80°C until use, and thawed only once. Protein quantification was carried out after thawing in a Synergy HTX Multi-Mode Reader (Biotek).

### Immunizations

To evaluate PD-L1 expression on plasmablast generated by TD-antigen immunization two different schedules were performed: One group of C57BL/6 mice were immunized intravenously with 2 × 10^8^ SRBCs (Applied Biological Products Management, Australia) and plasmablasts phenotype was analyzed at 7 days post immunization (DPI). Another group of C57BL/6 mice were transferred, by iv injection, with splenocytes from CD45.1 SWHel mice (Het/Het) containing 30.000 CD19^+^Hel^+^ cells simultaneously with 2×10^8^ Hel-conjugated SRBCs. Frequency of plasmablast and plasmablast phenotype was analyzed at 7 DPI.

### Bone Marrow Chimeras

Recipient Rag1−/− mice were sublethally irradiated with 500 Rad and reconstituted via iv injection with 10 × 10^6^ bone marrow-derived cells of the genotypes indicated below. Chimeric mice were maintained on antibiotics for 6 weeks after reconstitution and experiments were performed 8 to 10 weeks after reconstitution. Recipient mice were reconstituted with bone marrow from Blimp^fl/fl^-iCre^CD23^ (plasmablast deficient), or C57BL/6 (WT), or a combination of an 80:20 mixture of 10.10^6^ BM cells from Blimp^fl/fl^-iCre^CD23^ plus PD-L1 KO donor mice, respectively.

Chimera mice were infected with a previously tested dose of parasites, because chimeras presented an increased susceptibility to the infection. The experiments were performed at 14 Dpi following disease scores and ethical guidelines for mouse wellbeing.

### Cell Preparation

Spleens were obtained, and tissues were homogenized through a 0,70um cell strainer. Erythrocytes in cell suspensions were lysed for 5 min in Tris–ammonium chloride buffer. Viable cell numbers were determined by trypan blue exclusion using a hemocytometer.

### Flow cytometry

For surface staining, single-cell suspensions were washed first in ice-cold PBS and incubated for ten minutes with a Live/Dead Staining. Cells were washed in PBS and then incubated with fluorochrome labeled-Abs for 30 min at 4°C (surface staining). For intracellular cytokine staining, cells were cultured for 5 h with 50 ng/ml PMA (phorbol 12-myristate 13-acetate) (Sigma), 1,000 ng/ml ionomycin (Sigma) and Brefeldin A (eBioscience) or Monensin (eBioscience). Cells were fixed and permeabilized with BD Cytofix/Cytoperm and Perm/Wash (BD Biosciences) according the manufacturer’s instructions. Data were collected on a BD FACSCanto II, BD LSR II, or BD Fortessa X20, and were analyzed using the FlowJo software (TreeStar). The specific cell populations and gating strategy in each case are described in Supplementary Fig. 3.

### Antibodies

The following anti-mouse antibodies were used for FACS: CD19-PE (6D5), CD19-BV605 (6D5), CD3-AlexaFluor700 (17A2), CD3-APCCY7 (17A2), CD4-APC-Cy7 (GK1.5), CD8-PerCP-Cy5.5 (53-6.7), CD8-FITC (5H10-1), Streptavidin BV605, PD-1-BV421 (29F.1A12), PDL-1-Pecy7 (10F.9G2), Bcl6-PE-Dazzle594 (7D1), ICOS-AlexaFluor488 (C398.4A), CD45.1-BV605 (A20), CD11b-APCCy7 (M1/70), B220-APCCy7 (RA3-6B2), Live Dead Aqua 430 were from Biolegend; CD19-AlexaFluor700 (1D3), CD19-FITC (1D3), IgD-FITC (11-26c), CD8-PECy7 (53-6.7), TNF-APC (MP6-XT22), PD-L1-PE (MIH5) and CD-39-PerCP-eFluor710 (24DMS1) were from eBioscience; CD138-BV605 (281-2), CD138-APC (281-2), CD138-PE (281-2), CD138-Biotin, CD38-BV421 (90/CD38), CXCR5-Biotin (2G8), IFN-γ-FITC (XMG1.2), IL-4-PE (11B11), Ki-67-AlexaFluor647 (B56), Blimp-1-PE-CF594 (5E7), and CD8-BUV805 (53-6.7) were from BD Biosciences.

The following anti-mouse antibodies were used for tissue immunofluorescence: B220-PE (RA3-6B2) and CD3 FITC anti-CD3 (145-2C11) from Biolegend and CD138-APC (281-2) from BD Biosciences.

HEL was conjugated with AlexaFluor647 with a protein labeling kit (Invitrogen).

### Immunofluorescence

For tissue immunofluorescence, spleens from infected WT mice were collected and frozen over liquid nitrogen. Frozen sections 7 µm in thickness were cut, fixed for 10 min in cold acetone, left to dry at 25 °C and stored at −80 °C until use. Slides were hydrated in TRIS buffer and blocked for 30 min at 25 °C with 10% normal mouse serum in TRIS buffer. After blockade, slides were incubated for 50 min at 25 °C with the different Abs diluted in TRIS Buffer. Slices were mounted with FluorSave (Merck Millipore). Images were collected with an Olympus microscope (FV1000) and those recorded in the far-red channel were pseudo-colored blue.

### Cell culture

Splenic cells from infected mice were obtained at 14 Dpi and splenocytes were stained with fluorochrome-conjugated anti-CD19, anti-CD138 and anti-PD-L1. The elimination of plasmablasts, PD-L1^hi^, PD-L1^low^ and non-B cells from the splenocytes population was carried out by cell sorting in a FACSAria II (BD Biosciences) using different gates strategies (see Supplementary Fig.1).

Total splenocytes from infected mice or splenocytes depleted from plasmablast, non-B cells, PD-L1^hi^ and PD-L1^low^ cells were incubated with AgTpY (5ug/mL) plus anti-PD-1, anti-PD-L1, anti-PD-L2 (5ug/mL) or control antibodies (5ug/mL). After 48h of culture, TNF and IFN✉ were quantified by ELISA in the culture supernatant according to the manufacturer’s instructions (eBioscience).

### Statistics

Statistical significance through comparison of mean values was assessed by a two-tailed Student’st-test, one-way or two-way ANOVA followed by Bonferroni’s posttest using GraphPad software.

## ACKNOWLEDGEMENTS

We thank MP Abadie, MP Crespo, V Blanco, F Navarro and D Lutti from CIBICI-CONICET and A Cook and J Cappelo from JCSMR for their excellent technical assistance.

This work was supported by grants from the Agencia Nacional de Promoción Científica y Técnica (PICT 2015-0645), CONICET and R01AI116432. EAR, CLM and AG are Researchers from CONICET. MGS, FFV, CGB, MCR, JTB, and LA thank CONICET for the fellowship awarded. CGV was supported by Australian NH&MRC program, project and fellowship grants.

The authors declare no competing financial interests.

## AUTHOR CONTRIBUTIONS

MGS performed and designed most of the experiments, analyzed data and prepared figures and manuscript. FFV, CGB, MCR, JTB and LA collaborated with experiments performance. EIZ supervised design, analysis and interpretation of LCMV experiments. EW designed, performed, analyzed and interpreted LCMV experiments. IAC provided *P. chabaudi* parasites and contributed to the design of *Plasmodium* experiments, IAC and YC performed Plasmdium experiments.CLM and EVAR contributed to study design and analysis. AG and CGV conceived, designed and supervised the study and wrote the manuscript. All authors reviewed the manuscript before submission.

## ADDITIONAL INFORMATION

**Supplementary Information** accompanies this paper (Fig. S1–S3).

---

**Supplementary Figure 1.**
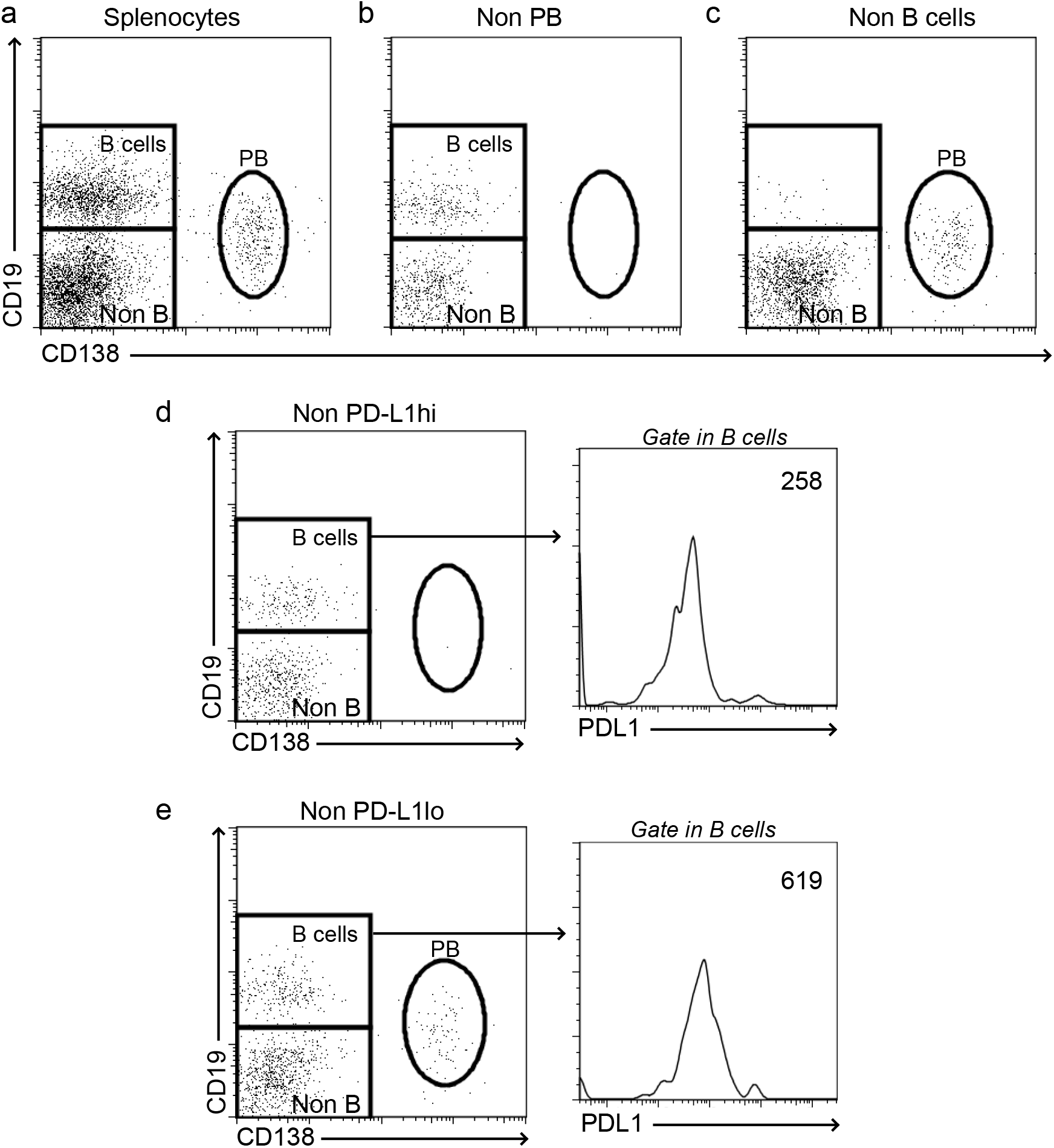
Gating strategies to determine the frequency of plasmablasts (PB), B cells and non-B cells (Non-B) in the spleen of C57BL6 mice infected with *T. cruzi* Y strain obtained at 14 Dpi. Splenocytes were stained with anti-CD19, -B220, -CD138 and anti-PD-L1 and depleted from plasmablasts (PB), non-B cells, and PD-L1^hi^ and PD-L1^low^ cells. Representative dot plot of: **a)** total splenocytes prior to the sort, **b)** post-sort excluding plasmablast (Non PB) and **c)** excluding B cells (Non-B cells). **d-e)** Dot plot representative of splenocytes without PD-L1^hi^ (Non PD-L1^hi^) and without PD-L1^low^ (Non PD-L1^low^) cells, respectively. The mean fluorescence intensity corresponding to PD-L1 expression on the remaining B cells is shown with the histogram.

**Supplementary Figure 2.**
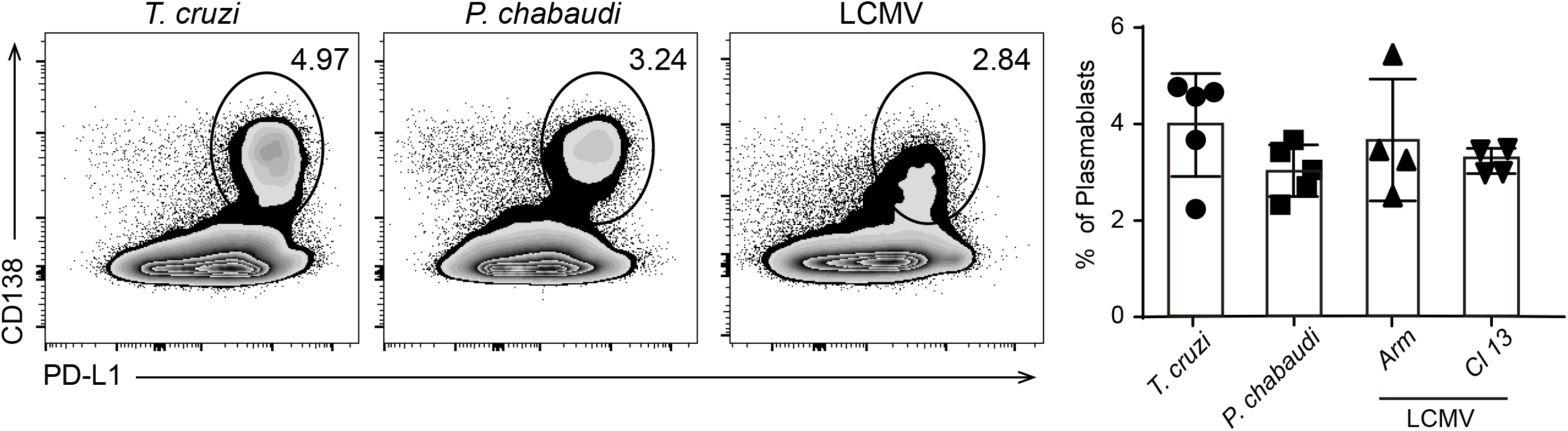
Representative flow cytometry plots and statistical analysis of the percentage of splenic PD-L1^+^ plasmablasts from *T. cruzi-, P. chabaudi-* or LCMV Cl13-infected mice. C57BL6 mice were infected with *T. cruzi* Y strain (n=5), *P chabaudi* (n=5) or with LCMV Cl13 or Arm (n=5) and the spleens were obtained at 14, 10 or 9 Dpi, respectively. Splenocytes were stained with anti-B220, anti-CD138 and anti-PD-L1.

**Supplementary Figure 3.**
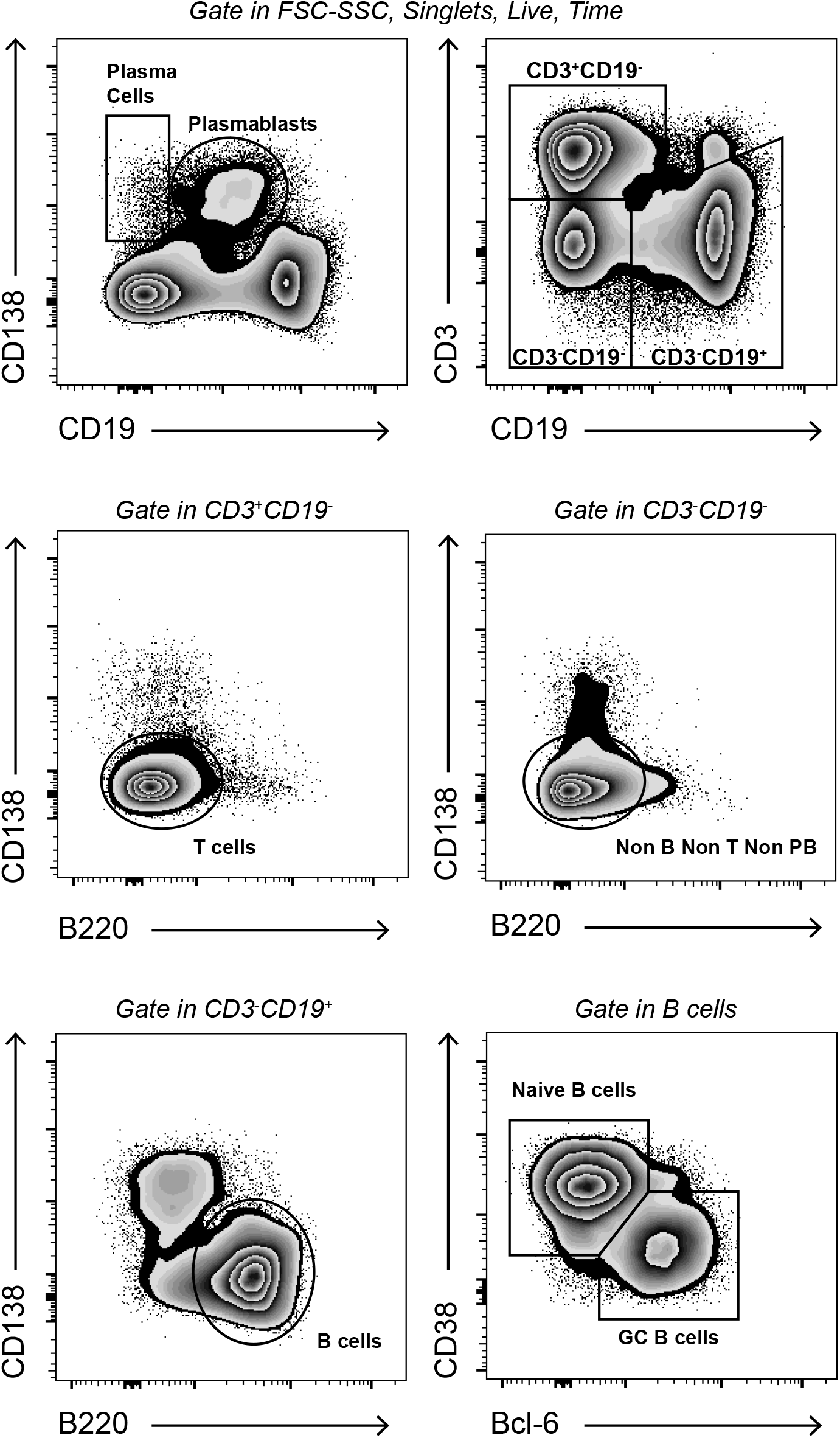
Gating strategies used for cell identification. Gating strategy to determine the percentage of plasma cells (CD19^−^CD138^+^) and plasmablasts (CD19^low^CD138^+^) and CD3^+^CD19^−^ (non B cells), CD138^−^CD3^−^CD19^+^ (B cells) and CD38^+^Bcl-6^neg^ CD3^−^CD19^+^ (naïve B cells) and CD38 ^neg^Bcl-6^+^ CD3^−^CD19^+^ (GC B cells) from the spleen of *T. cruzi* infected mice.

